# Modulation of auditory responses by visual inputs in the mouse auditory cortex

**DOI:** 10.1101/2021.01.22.427870

**Authors:** Sudha Sharma, Hemant Kumar Srivastava, Sharba Bandyopadhyay

## Abstract

So far, our understanding on the role of the auditory cortex (ACX) in processing visual information has been limited to infragranular layers of the ACX, which have been shown to respond to visual stimulation. Here, we investigate the neurons in supragranular layers of the mouse ACX using 2-photon calcium imaging. Contrary to previous reports, here we show that more than 20% of responding neurons in layer2/3 of the ACX respond to full-field visual stimulation. These responses occur by both excitation and hyperpolarization. The primary ACX (A1) has a greater proportion of visual responses by hyperpolarization compared to excitation likely driven by inhibitory neurons of the infragranular layers of the ACX rather than local layer 2/3 inhibitory neurons. Further, we found that more than 60% of neurons in the layer 2/3 of A1 are multisensory in nature. We also show the presence of multisensory neurons in close proximity to exclusive auditory neurons and that there is a reduction in the noise correlations of the recorded neurons during multisensory presentation. This is evidence in favour of deep and intricate visual influence over auditory processing. The results have strong implications for decoding visual influences over the early auditory cortical regions.

**Significance statement:** To understand, what features of our visual world are processed in the auditory cortex (ACX), understanding response properties of auditory cortical neurons to visual stimuli is important. Here, we show the presence of visual and multisensory responses in the supragranular layers of the ACX. Hyperpolarization to visual stimulation is more commonly observed in the primary ACX. Multisensory stimulation results in suppression of responses compared to unisensory stimulation and an overall decrease in noise correlation in the primary ACX. The close-knit architecture of these neurons with auditory specific neurons suggests the influence of non-auditory stimuli on the auditory processing.

## Introduction

Multisensory integration allows seamless assimilation of information from different sensory modalities to better perceive the environment. Cross-modal interaction of incoming sensory information is important since each sensory system is capable of encoding the same stimulus in a uniquely advantageous way(Maddox et al., 2015; Bizley et al., 2016). So far, most of our understanding of visual influence on the auditory pathway is with respect to sound source localization by subcortical regions(Werner-Reiss et al., 2003; Bulkin and Groh, 2012) and little is known of it in the ACX(Kayser et al., 2010; Atilgan et al., 2018; Morrill and Hasenstaub, 2018). However, other functions like communication and speech perception are higher cognitive functions(Ghazanfar, 2005, 2009; Skipper et al., 2007; Koelewijn et al., 2010; Romanski and Hwang, 2012) and a cross-talk between visual and auditory cortices is necessary for enhanced speech perception. Therefore, a basic understanding of this architecture in the animal models can help us decipher the framework over which these complex processes are built. Several studies have shown the cross-modal nature of sensory cortices(Ghazanfar and Schroeder, 2006). However, there is growing number of studies that deal with auditory influence over visual cortex(Ibrahim et al., 2016a; Murray et al., 2016; Meijer et al., 2017; Deneux et al., 2018; Knöpfel et al., 2019) rather than vice versa(Bizley and King, 2008a; King and Walker, 2012; Atilgan et al., 2018; Morrill and Hasenstaub, 2018). This is likely due to the easy accessibility of visual cortex with advanced tools like 2-photon imaging, which has not been used to explore the role of the mouse ACX in multisensory processing yet.

Although there is enough evidence of direct projections from the primary visual cortex to auditory areas in rodents other than mice(Budinger et al., 2006; Bizley et al., 2007; Campi et al., 2010), they are much less dense than those in the reverse direction(Oh et al., 2014; Ibrahim et al., 2016a). The reason for this disparity is unknown. Many physiological studies of visual influence over the A1 in ferrets and in primates have shown to affect the phase of oscillatory activity in the ACX, causing amplification of the response to an auditory stimulus(Ibrahim et al., 2016a; Atilgan et al., 2018) in presence of an audio-visual stimulus. This amplification leads to decreased response latencies and enhanced responses in a noisy background(Wang et al., 2008). Other studies also report spiking activity in the ACX by visual stimuli, but also that this spiking is limited to infragranular layers(Bizley et al., 2007; Morrill and Hasenstaub, 2018) and absent in supragranular layers. These studies have resorted to electrophysiology, which lacks spatial resolution. This has led to the assumption that the visual responses in the ACX either are the result of local field potential modulation from the surrounding visual cortex or are only present in the deeper layers of the auditory.

Using two-photon calcium imaging with GCaMP6s in awake mouse, we probed layer 2/3 (supragranular) of the ACX with auditory, visual, and audio-visual (or multisensory) stimuli. In addition to responses to auditory stimulation (broadband noise), we found robust Ca2+ based responses to visual stimulation in layer 2/3 of primary as well as secondary ACX. Most of the neurons show suppressive responses to multisensory stimulus as compared to their unimodal counterparts. This study also reveals neurons having different response strengths towards the auditory, visual and multisensory stimuli. Two-photon calcium imaging also reveals the close-knit spatial architecture of the auditory and multisensory neurons, as well as an effect of multisensory stimuli on the noise correlations in the primary as well as secondary auditory areas. This paper is an attempt towards understanding the complex nature of multisensory processing in the ACX by studying the inherent capabilities of these neurons to perform multisensory processing without undergoing any plasticity due to multisensory experiences.

## Materials and Methods

All procedures were in accordance with protocols approved by the Indian Institute of Technology Kharagpur, Institutional Animal Care and Use Committee and guidelines of the National Institutes of Health. Mice were acquired from Jackson Laboratories (PV-Cre [JAX 008069], SOM-IRES-Cre [JAX 013044], ROSA LSL-tdTomato [JAX 007908]) housed in a room with a reversed light cycle. Experiments were performed during the dark period.

### Surgery

Adult mice (>30 days old, male and female) were anesthetized with isoflurane (5% induction and 1% maintenance), injected with dexamethasone (2 mg/kg body weight) intraperitoneally and placed on a stereotaxic frame. Body temperature of the animal was maintained at 37 °C throughout the procedure using a heating pad. Incision was made to expose the skull. A craniotomy (3mm diameter) was done to expose the ACX. Virus (AAV.Syn.GCaMP6s.WPRE.SV40) was loaded into a glass micropipette mounted on a Nanoject II attached to a micromanipulator and injected at a speed of 20 nL per min at several places exposed by the craniotomy in PV-tdTomato and SOM-tdTomato transgenic animals. The craniotomy was covered with a glass coverslip(Goldey et al., 2014). Mice were left in their home cage to recover and virus to integrate and express for 2 weeks. Experiments were conducted 15-20 days after viral injection.

### Histology

Brains from all mice used for recording were harvested post hoc all experiments to confirm recording locations. Mice were deeply anesthetized with isoflurane followed by transcardial perfusion with 0.1M phosphate-buffered saline followed by 4% paraformaldehyde. Brains were harvested and after a post-fixation period of 8-10 hours in 4 °C, 100 um thick sections were cut using a vibratome (Leica VT1000S) and images were taken in a fluorescence microscope (Leica DM2500).

### Auditory and Visual stimulation

Auditory stimuli were presented to the ear contralateral to the recording hemisphere inside a light-proof chamber, 10 cm away from the right ear (contralateral) of the mouse. Stimulus was generated through custom-written software in MATLAB (Mathworks), passed through TDT RZ6 multifunctional processor. Acoustic calibrations were performed using microphone (Brüel & Kjær, Denmark) and showed a typical flat (+/-7 dB) calibration curve from 4-60 kHz. All mice were first presented with pure tone (*T* stimulus, 5 repetitions each, 6-48 kHz, 1/2 octaves apart, 50ms duration with 5ms rise and fall, 5s inter stimulus interval at ∼65-75 dB SPL) in order to identify the tonotopic axis. For comparison of unisensory vs. multisensory responses, we chose to play white noise (*WN*, bandwidth 6-48 kHz, 10 repetition) instead of pure tones as the auditory stimulus (to be later paired with visual LED flash for multisensory stimulation) in order to avoid the influence of frequency selectivity of individual neurons. For visual stimulation, a white LED (5 mm Round White LED(T-1 3/4)) kept 10 cm away from the eye of the animal, close to the source of sound, was used. Full-field illumination to the contralateral eye was provided and the intensity (1-160 lux, Model: LX1010b, Digital Lux Meter, 0 to 20000 lux) was controlled through the NI DAQ card, using MATLAB routines. LED blink stimulus was presented for 10 ms (*V* stimulus) with an interstimulus interval of 5s in order for the cones in the retina to recover. Audio-visual stimulation (or Multisensory, *M* stimulus) comprised of simultaneous presentation of the *A* and *V* stimuli (*M*), as above. 10 repetitions of the *M* stimuli were presented.

### Widefield calcium imaging

Images were acquired using a 14-bit CCD CoolSNAP HQ2 CCD camera (Photometrics). Images of surface vasculature were acquired using blue LED illumination (470 nm), and wide-field GCaMP signals were recorded (36 ms frame period) using blue illumination (470 nm). Fluorescence signals were acquired at 4x binning of 1040×1392 (260×348) pixels using a 4x objective(Olympus). Each trial consisted of 50 baseline frames and 50 frames of response. A 500 ms sound stimulus (60-30dB SPL pure tone with a frequency in the range 4-48 kHz, 30 trials for each frequency) was presented starting at the 51^st^ frame. The inter-trial interval was 5-7 s. Images during the response period (0.5–2 s from the sound onset) were averaged, and the average image during the baseline was subtracted from it. This difference was divided by the average image during the baseline. 200-300 ms after stimulus start was considered as a response window.

### 2-photon calcium imaging

Two-photon calcium imaging was performed using a commercially available 2-photon microscopy system (Prairie View Technologies). The microscope was controlled by Prairie View software (Prairie View Technologies). For imaging, the excitation beam of wavelength 860-920nm generated from Insight laser (Spectra-Physics, USA) was directed over the glass-covered brain surface in layer 2/3 (180-250um depth). The laser was delivered through 20X/0.8 NA water immersion objective (2mm WD, Olympus). The laser power was adjusted from 50mW to 80mW depending upon the condition of the specimen. Frames in the region of interests (∼150 μm x 200 μm with 1.16 μm pixel size) were imaged at ∼4-6 Hz (160-250ms frame period, 1.2-4 μs dwell time) with the stimulus presented at 10th frame in a sequence of 20 frames per stimulus. Cross-correlation-based image alignment was used for motion correction in XY plane. Center of cells was marked manually and rings were drawn within of 5µm radius around the point of selection. Luminance(F_l_) of all the pixels within the rings was the mean of all pixel values within the marked region. Baseline fluorescence was subtracted to calculate change in fluorescence and normalized by the baseline fluorescence to compute dF/F. All other analysis was done on these dF/F values.

### Data Analysis

#### Widefield Imaging data analysis

For creating a tonotopic map, we used 5^th^ to 10^th^ frame (180-360 ms) after the stimulus as the response frames. Mean along the 30 iterations was calculated for each of the response frames. Using these mean responses frames, we computed the dF/F by subtracting the mean of baseline frames and then dividing by it. The dF/F was smoothed by a gaussian filter with standard deviation of 2. Next, this dF/F matrix was normalized by the maximum to bring the range between -1 and 1. To this matrix, we applied a threshold above which all values were set to 1 and pixel values below threshold were set to zero. The threshold used was generally 65-75 % of the maximum pixel value. The three image matrices, one each for low, middle and high frequencies were overlaid on each other to illustrate the tonotopic axis (Fig. 1, top row, right).

**Figure 1.**
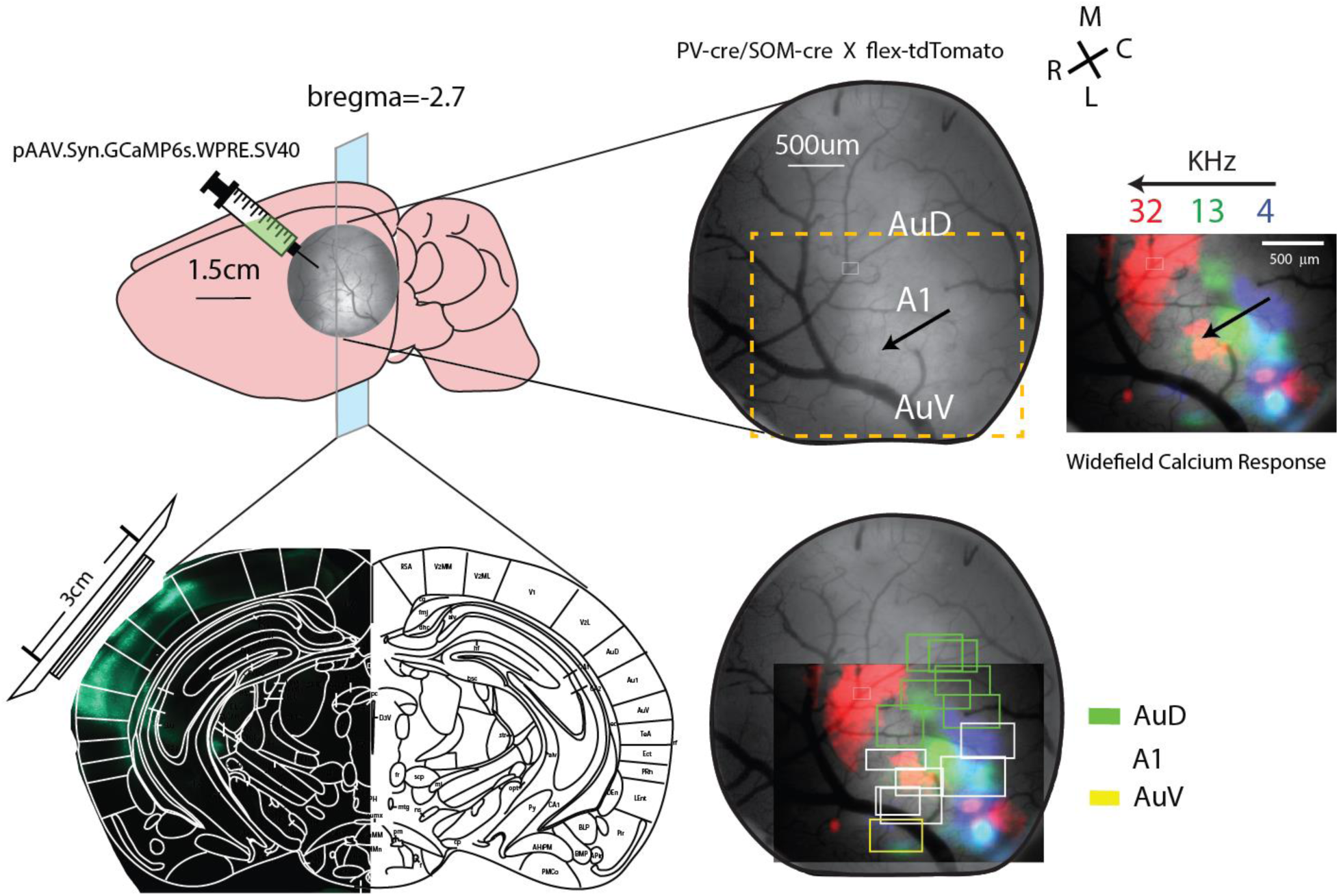
Demarcation of areas based on tonotopic organization obtained via Widefield Calcium imaging responses to different frequencies. **Top left**, cartoon showing mouse brain and placement of glass window for long term calcium imaging. **Bottom left**, cross section of mouse brain showing medio-lateral extent of the injection sites in the ACX. **Top right**, example image of the cortical surface obtained through the glass window under a blue fluorescent light source. On the right of it is the tonotopic map of frequencies showing rostro-caudal extent of A1 obtained via widefield calcium imaging. **Bottom right**, small boxes in different colours represent regions imaged via 2 photon calcium imaging over days and the auditory region they belong to. Green: AUD, White: A1, Yellow: AuV.

#### Demarcation of auditory areas (Ventral ACX (AuV), A1,Dorsal ACX (AuD))

Recordings were done on the ACX in five animals injected with AAV.Syn.GCaMP6s in the ACX. All animals were exposed to low, middle and high frequencies for identifying the A1 tonotopic axis and to mark A1 within the craniotomy and an LED flash for demarcating the visual cortex boundary medial to AuD based on wide field calcium responses to LED. Based on the responses obtained, the imaging region was divided into four sub regions, secondary visual cortex (V2) which showed highest response activity to visual stimulus, A1 containing the tonotopic axis, AuD is the region between V2 and A1, and AuV to be below the marked A1 boundary. All recorded ROIs were then assigned a region, depending on their location. For this study, we excluded ROIs from the V2 region from our analysis.

#### Calculation of significant response and its reliability

To identify a significant response to a given stimulus, we compared the mean baseline activity of three frames before the stimulation with the mean response activity of three frames after stimulation in all iterations using one-tailed paired t-test. We considered *p* value less than 5% for the response by excitation and less than 10% for the response by hyperpolarization as our criterion for response. We also calculated the reliability of responses by comparing the mean signal of the response frames in iteration with the mean of baseline frames. If this value was greater than 1.5 standard deviation for excitation (and 1 standard deviation for hyperpolarization) of the baseline signal, it was considered as a response in that iteration. Reliability was calculated as the ratio of number of responding iterations and total iterations. Keeping the standard deviation to calculate reliability and the criterion for significance same for both type of responses, we were unable to get the hyperpolarization responses because the calcium indicators cannot effectively capture hyperpolarization(Chamberland et al., 2017).

#### Classification of neurons

We divided the neurons into different classes based on the stimulus they responded to. Neurons that responded only to visual stimulus are *V0*, neurons that responded only to auditory stimulus are *A0*, neurons that responded only to multisensory stimulus are *M0. N*eurons that responded only to auditory stimulus but showed significant changes in response on presentation of multisensory stimulus are multisensory neurons type *MA*, neurons that responded only to visual stimulus but showed significant change in response on presentation of multisensory stimulus are multisensory neurons type *MV. N*eurons that responded to auditory stimulus and visual stimulus irrespective of response to multisensory stimulus are multisensory neurons type *MAV*.

| Cell types | V | A | MS |
| --- | --- | --- | --- |
| VO | 1 | 0 | 1 |
| MV | 1 | 0 | 1* |
| MO | 0 | 0 | 1 |
| MAV | 1 | 1 | 0/1 |
| AO | 0 | 1 | 1 |
| MA | 0 | 1 | 1* |

\* significantly different than the unisensory counterpart

### Bootstrap method for significance

a. **Comparisons of reliabilities:** For comparison of reliabilities for a given population between different stimuli, bootstrap method (Efron and Tibshirani, 1993) was performed to obtain confidence intervals on the reliabilities. Bootstrap populations (*n* = 1000) were created by randomly sampling from reliabilities with replacement from the original dataset. We obtained 1000 reliability values from the population of 1000 bootstrap sets. The mean reliabilities from two different conditions were considered to be significantly different if the 99% confidence intervals of the two did not overlap with each other and accordingly for one-sided comparisons.
b. **Spatial organization of neurons w.r.t type A0 neurons:** Distance between type A0 and its nearest type x (where, x= V0/MV/MA/MAV/A0) was obtained. These distances were averaged to obtain the mean distance (raw) between type A0 and type x neurons. Bootstrap population (*n*=1000) was created by randomly sampling with replacement (200 times) from these distances and averaged to obtain the confidence intervals from the mean values for comparison between populations. The raw mean distances from two different pairs were considered significantly different if the 95% confidence intervals of the two did not overlap with each other. We used a similar procedure to obtain the significance levels in the mean probability values within 50um region around type A0 neurons with a criterion of at least five cells to be present within 50um region.

## Results

### Cross-modal influence is more prevalent in neurons of the ACX than their unisensory nature

2-photon Ca^+2^ imaging was performed in layer 2/3 of awake mice (3 animals (SOMxtd-tomato) and 2 animal (PARVxtd-tomato)) after implanting a cranial window with injections of AAV-GCaMP6s targeted at layer 2/3 in multiple regions of the ACX. Using tonotopic axis as reference, ROIs were classified to belong to a particular auditory field (Fig. 1, Methods). For unisensory stimuli either Noise (50ms width, auditory) or a white LED flash (10 ms width, visual) was used. For multisensory stimulus the LED flash and noise were presented together with the same starting time and durations as mentioned above. Data from multiple ROIs in the ACX were collected over days. Each responding neuron was categorized into one of the six categories based on its response to different stimuli and region it belonged to as shown in Table 1 (Methods, *Classification of Neurons*).

**Table 1.** Number of responding neurons in each category in different areas. S, suppression; E, enhancement.

|  | V0 | MV(S,E) | MAV(S,E) | A0 | MA(S,E) | M0(S,E) | TOTAL |
| --- | --- | --- | --- | --- | --- | --- | --- |
| **AUD** | 55 | 257(195,62) | 108(17,8) | 215 | 372(335,37) | 328(23,305) | 1335 |
| **A1** | 32 | 160(156,4) | 359(73,5) | 842 | 847(830,17) | 249(65,184) | 2489 |
| **AUV** | 58 | 119(107,12) | 299(26,11) | 167 | 186(173,13) | 266(8,258) | 1095 |
| **TOTAL** | 145 | 536(458,78) | 766(116,24) | 1224 | 1405(1338,67) | 843(96,747) | 4919 |

Contrary to previous reports(Morrill and Hasenstaub, 2018), layer 2/3 of all regions of the ACX was found to have neurons responding to visual stimulus, including neurons that responded only to visual stimulus (type V0) and not to auditory stimulus (Fig. 2A, Top row(left), Table 1). AuD and AuV had a greater proportion of V0 type neurons compared to A1 (AuD (4.1%, 55/1335), A1(1.2%, 32/2489), AuV (5.2%, 58/1095)) (see Fig. 2B, Left and Table 1) while the greatest proportion of type A0 neurons were present in A1(AuD (16.1%, 215/1335), A1(33%,842/2489), AuV (15.2%, 167/1095) (Fig. 2B, middle). Multisensory neurons were prevalent in all auditory regions. However, there was some specificity observed with respect to the subtype of multisensory neurons and the region involved. *MV* type neurons were maximum in AuD (19.2%, 257/1335), followed by AuV (10.8%, 119/1095) and A1(6.4%, 160/2489) (Fig. 2B, right), while *MA* type neurons were present with greatest proportion in A1 (34%, 847/2489) and less in AuD (27.8%, 372/1335) and even lesser in AuV (16.9%, 186/1095) (Fig. 2B). *MAV* type neurons were mostly present in AuV (27.3%, 299/1095) with highest proportion followed by A1 (14.4%, 359/2489) and AuD (8.0%, 108/1335). Apart from these, M0 formed a significant proportion of population responsive exclusively to multisensory stimulus and not responsive to the presentation of individual unisensory stimulus in all three regions (AuD=24.5%, 328/1335, A1=10.0%, 249/2489, AuV=24.2%, 266/1095). Overall, more than 60% of cells in all three regions were multisensory in nature either responsive to only multisensory stimulus (type M0) or to both unisensory stimuli (type MAV) or undergoing cross modal modulation (type MA and MV). To discern the role of PV+ and SOM+ inhibitory interneurons in multisensory processing, we looked at the distribution of these neurons into different categories. We did not observe anything unusual that could point to their particular role. In fact, the distribution of PV+ and SOM+ neurons into the categories looked similar in proportions to the one in Fig. 2B. We did not explore their function further.

**Figure 2.**
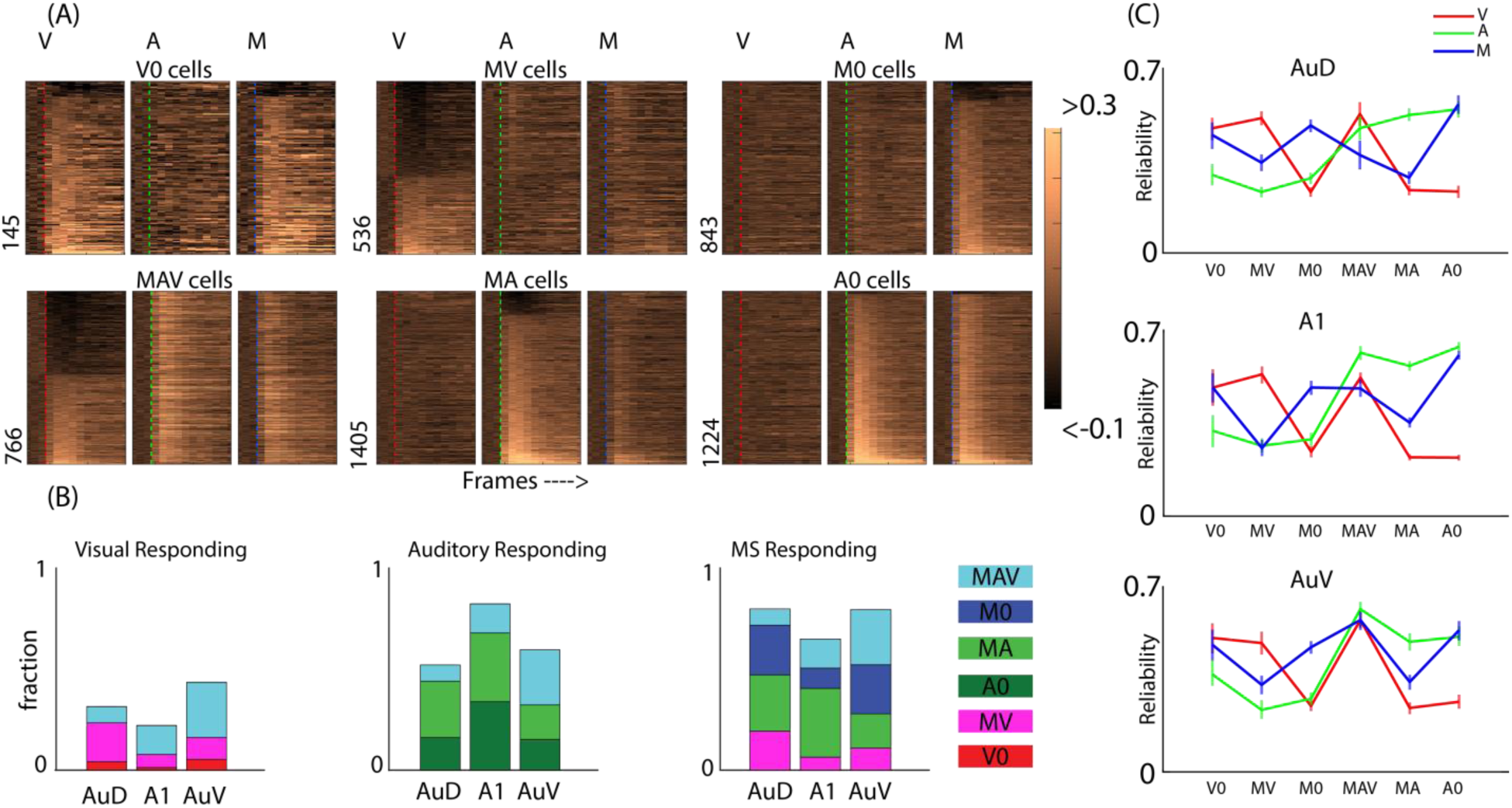
Classification, proportion and reliabilities of all cell types in different regions of the ACX. **A**, Top row(left), type V0 cells with their mean df/f values for auditory(A), visual(V) and multisensory(M) stimulation over total recorded frames. Red, Green and Blue lines on the images shows stimulation frame. Same scheme is repeated for type MV, M0 cells in the top row and MAV, MA and A0 cells in the bottom row. **B**, Proportion of neuron types, indicated by colour, as a fraction of total responding neurons in a region, Red: type V0 neurons, Pink: type MV neurons, Dark green: type A0 neurons, Light green: type MA neurons, Blue: type M0 neurons, Cyan: type MAV neurons. **C**, Mean reliability values with confidence intervals obtained from bootstrap analysis of each type of population for region AuD(top), A1(middle), AuV(bottom). Green: mean reliabilities on auditory stimulation, red: mean reliabilities on visual stimulation, blue: mean reliabilities on multisensory stimulation.

We also looked at the mean reliability of response of the six classified neuron types to different stimuli. Type A0 and type MA neurons in A1 have high response reliability for auditory stimulus and low for visual stimulus (one-sided comparison of mean reliabilities based on bootstrap confidence intervals, A0(A)=0.63[0.62, 0.65], A0(V)=0.22[0.21, 0.23], MA(A)=0.56[0.55, 0.58], MA(V)=0.22[0.21, 0.23]). While type V0 and type MV neurons have higher response reliability for visual stimulus and low for auditory stimulus (V0(V)=0.46[0.41, 0.55], V0(A)=0.28[0.26, 0.39], MV(V)=0.53[0.5, 0.57], MV(A)=0.27[0.24, 0.3], Fig. 2C). Type A0 and type V0 neurons show similar response reliabilities for multisensory stimulus as that to auditory and visual stimulus respectively (A0(A)=0.63[0.62, 0.65], A0(M)=0.60[0.59, 0.62], V0(V)=0.46[0.41, 0.55], V0(M)=0.49[0.43, 0.55], Fig. 2C). However, type MV and type MA neurons shows significant reduction in reliability to multisensory stimulation compared to visual and auditory stimulation respectively(MV(V)=0.53[0.5, 0.57], MV(M)=0.26[0.23, 0.29], MA(A)=0.56[0.55, 0.58], MA(M)=0.35[0.33, 0.37]). This is likely due to suppression of responses to multisensory stimulation which will be discussed later in the article. Type M0 neurons have higher response reliabilities for multisensory stimulation while type MAV neurons have high response reliabilities for all three stimuli (M0=0.24(V), 0.29(A), 0.48(M), MAV=0.52(V), 0.61(A), 0.48(M)). AuD and AuV also follow the same trend (Fig.2C). Responses for auditory stimulus in type A0 population were found to be more reliable in A1 compared to AuD and AuV (AuD=0.54 [0.51, 0.57], A1= 0.63[0.62, 0.65], AuV= 0.51[0.48, 0.54]). The distribution of response reliabilities is in agreement with our neuron classification scheme and shows the difference between different populations.

### Spatial organization of auditory specific and nonspecific neurons around each other

Contrary to expectation, multisensory neurons (MA, MV, MAV, M0) outnumber the type A0 neurons in all regions of the ACX (see Table 1 & example ROI in Fig. 3A). Given the high number of mutisensory neurons in the ACX, we looked at the local spatial organization of the primarily auditory type A0 neurons and multisensory neurons with respect to each other in the primary and secondary auditory fields. We chose type A0 neurons because they are specific to auditory stimulation against all other, grouped in type M (V0, MV, MAV, MA, M0), which are nonspecific to auditory stimulation. Assuming density of neurons in layer2/3 of all auditory regions to be similar, we compared the mean nearest neighbour distances between type A0 neurons and all other types, within and between auditory regions (Fig. 3B, Table 2). All comparisons were made based on the confidence intervals from the bootstrapped mean nearest neighbour distances (Methods, Table 2). We found that type A0 neuron is more likely to have another type A0 neuron as its nearest neighbour (AuD=34.86um, A1=21.03um, AuV=31.05um) followed by type MA (AuD=39.53um, A1=29.48um, AuV=45.69um) and type MAV in close proximity (AuD=53.21um, A1=39.64um, AuV=46.36) in all three regions (Fig. 3B, Table 2). Type M0 neurons are closer to type A0 neurons in AuD and AuV while they lie farther away in A1 (AuD=40.59um, A1=83.77um, AuV=37.29um, Fig. 3B). Type V0 lie furthest away in A1 while they are relatively closer in AuD and even closer in AuV (AuD=102.00um, A1=132.93um, AuV=89.60um). Type MV neurons lie significantly closer to A0 neurons than are V0 neurons (AuD=81.55um, A1=97.79um, AuV=68.04um).

**Table 2.** Mean nearest neighbour distance of a cell type from type A0 neuron with 95% confidence intervals with lower and upper bounds.

|  | AuD (mean [L, U])(in um) | A1 (mean [L, U]) (in um) | AuV (mean [L, U]) (in um) |
| --- | --- | --- | --- |
| **V0** | 102.00[92.19,111.58] | 132.93[121.37,145.00] | 89.60[78.10,101.03] |
| **MA** | 39.53[35.95,43.19] | 29.48[26.58,32.36] | 45.69[40.98,50.82] |
| **MAV** | 53.21[48.11,58.63] | 39.64[35.58,44.19] | 46.36[41.26,51.27] |
| **MV** | 81.55[74.79,87.85] | 97.79[89.64,106.15] | 68.04[62.65,73.63] |
| **M0** | 40.59[36.62,45.09] | 83.77[74.22,93.64] | 37.29[32.98,42.44] |
| **A0** | 34.86[30.21,39.49] | 21.03[18.83,23.61] | 31.05[28.13,34.39] |

**Figure 3.**
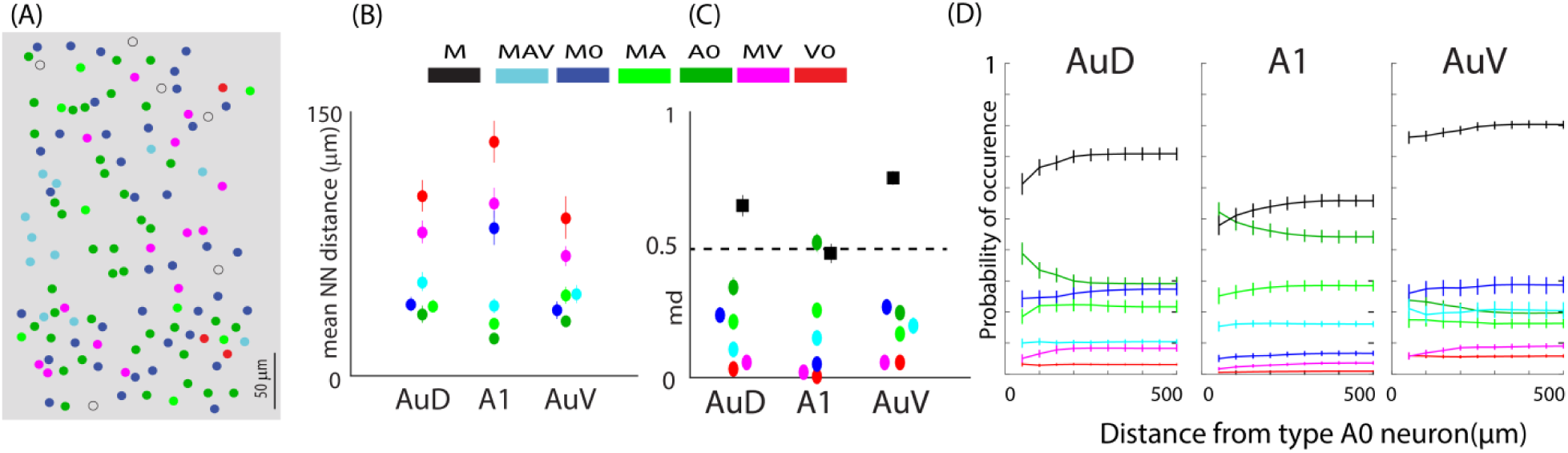
Spatial organization of different cell types w.r.t. type A0 neurons in a local region. **A**, Example recorded region with colour coded cell types. **B**, mean nearest neighbour (NN) distance between type A0 neuron and the nearest neuron of a particular cell type (colour coded) in AuD, A1 and AuV regions of the ACX. The coloured points are displaced to show the confidence intervals (also see table 2). **C**, mean probability of occurrence of a certain cell type (colour coded) within 50um radius around type A0 neuron in AuD, A1 and AuV regions of the ACX. The coloured points are displaced to show the confidence intervals (also see table 3)). **D**, Probability of occurrence of different cell types (colour coded) w.r.t type A0 within a given radii (x axis). Black: type M neurons, Cyan: type MAV, Blue: type M0, Light green: type MA, Dark green: type A0, Pink: type MV, Red: type V0 neurons.

We further looked at the probability of each type of neuron within 50um radius around type A0 neuron. If most of the neurons in a 50um radius around a type A0 neuron are of type A0, probability of its occurrence would be high, leading to auditory dominance in this region, and, if most of the neurons are of type M, there would be multisensory dominance (Fig. 3C). We looked at contributions made by each population type to the probability of type M individually as well (Figure. 3C, Colour coded). All comparisons were made on the mean probability values with 95% confidence intervals from the bootstrapped data (Table 3). The mean probability value of type M neuron around a type A0 neuron in A1 is 0.48 (Fig 3C, Table 3). Hence, within a radius of 50um around a type A0 neuron ∼50% of cells on average were found to be of type M while the other 50% are of type A0. While AuD and AuV are multisensory dominant regions (probability in AuD=0.65, AuV=0.75). If we look at the contribution of individual neurons types to the overall proabability value, type V0(AuD=0.03, A1=0.0, AuV=0.05) and type MV (AuD=0.05, A1=0.01, AuV=0.05) neuron are found very rarely within 50um in AuD and AuV while being completely absent in A1. Type MA makes a higher contribution (AuD=0.21, A1=0.25, AuV=0.16), followed by MAV (AuD=0.1, A1=0.15, AuV=0.19) in all three regions. Another difference lies in the type M0 population which has the highest contribution in AuD and AuV but not so much in A1 (AuD=0.24, A1=0.05, AuV=0.26).

**Table 3.** Mean probabilities of occurrence of a cell type within 50um radius of type A0 neurons with 95% confidence intervals with lower and upper bounds.

|  | AuD (mean [L, U]) | A1 (mean [L, U]) | AuV (mean [L, U]) |
| --- | --- | --- | --- |
| **M** | 0.65[0.61,0.69] | 0.48[0.44,0.51] | 0.75[0.72,0.77] |
| **VO** | 0.03[0.02,0.04] | 0.00[0.00,0.00] | 0.06[0.04,0.06] |
| **MA** | 0.21[0.18,0.24] | 0.26[0.22,0.28] | 0.16[0.13,0.19] |
| **MAV** | 0.10[0.08,0.12] | 0.15[0.13,0.17] | 0.20[0.16,0.23] |
| **MV** | 0.06[0.04,0.07] | 0.02[0.01,0.02] | 0.06[0.04,0.07] |
| **MO** | 0.24[0.21,0.27] | 0.05[0.03,0.06] | 0.27[0.23,0.29] |
| **A0** | 0.35[0.30,0.38] | 0.52[0.48,0.55] | 0.25[0.22,0.27] |

From these observations, we propose presence of multisensory processing hub in auditory cortical areas at the center of which is a type A0 neuron surrounded by MA and MAV in close proximity. Presence of type MAV neurons in close proximity of type A0 neurons needs special mention as it reflects a direct visual influence over auditory processing. Probability of occurrence of different cell types within a given distance from type A0 neurons in A1 (Fig. 3D, middle) shows a decrease in probability of occurrence of type A0 neurons around the same type (shown in dark green) and an increase in the type M neurons with distance (shown in black) in all three regions. Probability of occurrence of type M neurons remains high along with distance observed from type A0 neurons in secondary areas, while in A1, probability of occurrence of type A0 is high in the nearby regions and reverses after 150um(Fig. 3D, middle).

### Layer 2/3 neurons in A1 respond primarily by hyperpolarization to visual stimulus

22% of responding neurons in A1 respond to visual stimulation (V0, MV, MAV, see Table 1, example ROI in Fig. 4A&B). Majority of these neurons in A1 respond by hyperpolarization to visual stimulus (68%). Some fraction of neurons in secondary auditory areas also respond by hyperpolarization but majority of them respond by excitation (Fig. 4A, lower panel). Response by excitation could be a result of direct or indirect excitatory drive from the visual cortex, while the response by hyperpolarization must involve inhibitory neurons. In agreement with the above statement, we found several neurons which responded by hyperpolarization to visual stimulus and by excitation to auditory stimulus (Fig. 4C). In order to look at the role of inhibitory neurons in the processing of cross modal information, we looked at a specific population of A1 neurons, a subpopulation of type MAV, which respond by hyperpolarization to visual stimulus and by excitation to auditory stimulus which are 44% of all visual responding neurons in A1. For such a differential response to happen there has to be an inhibitory input on these neurons that respond to visual stimulus and does not respond to auditory stimulus (Fig. 4C upper panel). We looked for all the inhibitory neurons in the vicinity of these neurons which show such a behaviour but we found only two inhibitory neurons that show such a behaviour which is not enough to explain the observations.

**Figure 4.**
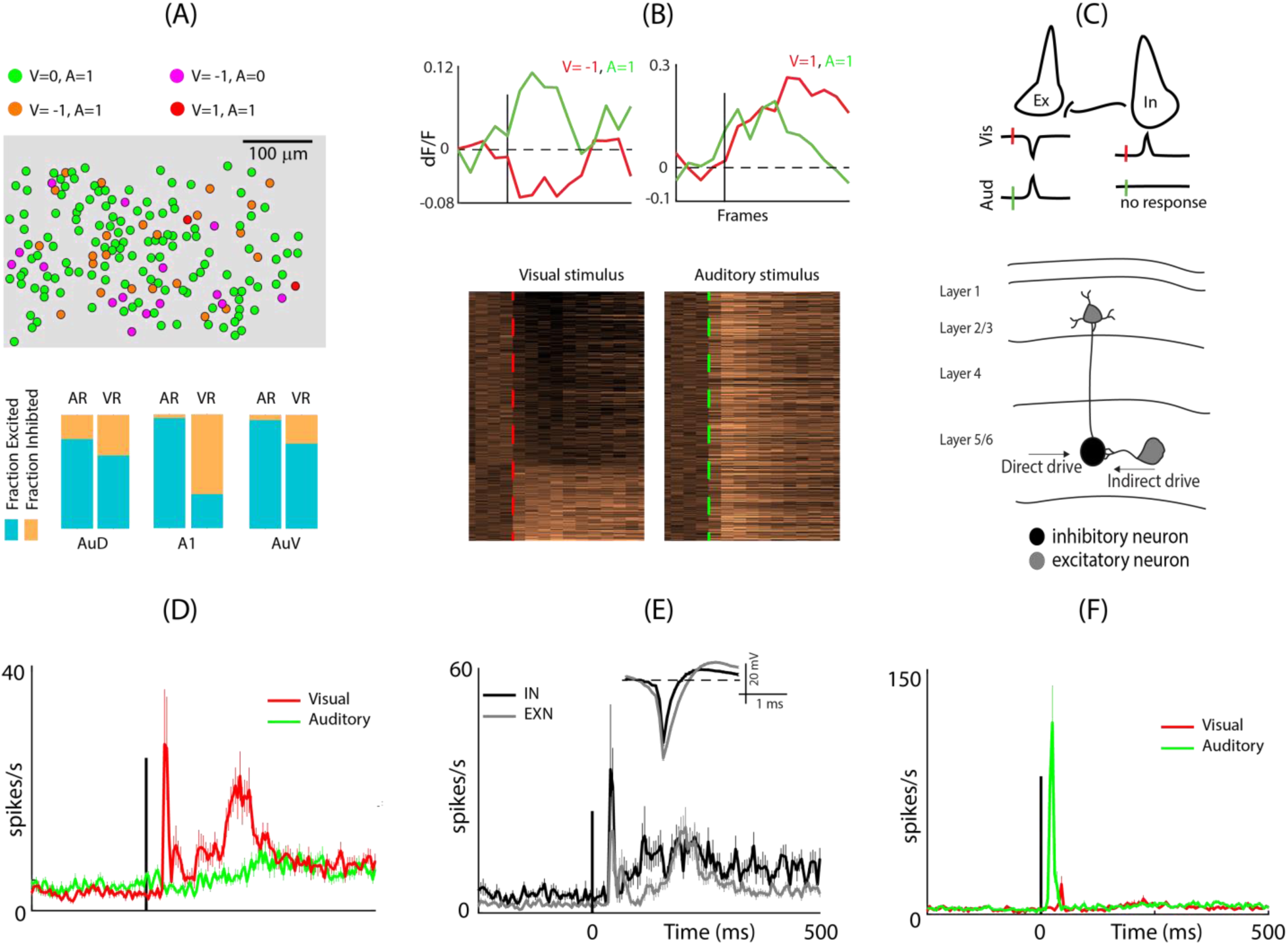
Neural underpinning of hyperpolarization behaviour of visual responding neurons in A1. **A (upper panel)**, Example ROI showing visual responding neurons in A1, colour coded by type of visual and auditory response. **A (lower panel)**, Fraction of cells showing excitation and inhibition for auditory stimulus (AR) and visual stimulus (VR) in different regions of the ACX. Blue, fraction showing excitation; orange, fraction showing hyperpolarization. **B(upper panel)**, Example traces of dF/F signal from the ROI in A (upper panel). **B (lower panel)**, response profiles of all visual responding cells in A1 for visual stimulus (left) and their response for auditory stimulus(right). **C (upper panel)**, cartoon diagram explaining the possible cause of hyperpolarization responses. **C(lower panel)**, hypothetical circuit model explaining the role of layer 5/6 inhibitory neurons in causing hyperpolarization of layer 2/3 neurons in A1. **D**, mean population response with SE of V1A0 neurons in layer 5/6 of the ACX. **E**, mean population response with SE of V1A0 inhibitory and excitatory neurons separated by color in layer 5/6 of ACX for visual stimulus with their respective spike shapes in the inset. **F**, mean population response of neurons with SE located near to the above mentioned (in D) neurons showing response to auditory and visual stimulation.

Thus, it is clear that the PV+, SOM+ inhibitory neurons of Layer 2/3 are not directly involved in carrying out differential response to visual and auditory stimuli. Therefore, following possibilities exist that can explain the response by hyperpolarization to visual stimuli and excitation to auditory stimuli. First, inhibition of layer2/3 neurons could be mediated by the inhibitory drive from Layer 1, which like in visual cortical areas has been shown to modulate responses by auditory stimulus (Ibrahim et al., 2016b; Chou et al., 2020). Second, this hyperpolarization could also be mediated by the excitation of layer5/6 inhibitory neurons either directly or indirectly (via another excitatory neuron in the ACX) by excitatory drive from visual cortex (Fig. 4C, lower panel). The axon terminals of these inhibitory neurons in layer 5/6 can potentially hyperpolarize the excitatory neurons in Layer2/3 of A1(Iurilli et al., 2012).

We recorded responses to visual and auditory stimulus from layer 5/6 of the ACX using a 4 by 4 array of electrodes (data also published in the paper (Sharma and Bandyopadhyay, 2020)). We separated the recorded neurons into inhibitory and excitatory neurons based on the spike shapes (Fig. 4E). We calculated the spike width, i.e. full width-half maximum followed by a threshold at 0.25 ms to separate the inhibitory from the excitatory neurons. We were able to identify a sub-population of units (48 units, 8% of all responding units) that did not respond to auditory stimulus but responded to a visual stimulus (V1A0_Ephys_, Fig. 4D), out of which 33 were excitatory and 15 were inhibitory neurons. To make sure that all our electrodes were in the ACX and not the nearby V2 area, which is expected to have plenty of such inhibitory and excitatory neurons, we looked at the auditory responses of close by units on the neighbouring electrodes (Fig. 4F). We found that most units responded robustly to the auditory stimulus and of these, ∼73% did not respond to the visual stimulus. Based on the latency of peak response to visual and auditory stimulus and the post-hoc Nissl staining of the sliced brain (Sharma and Bandyopadhyay, 2020), we claim that all of our electrodes were in the ACX. Thus, we claim the existence of inhibitory and excitatory neurons in layer 5/6 of the ACX as potential direct and indirect source that may drive hyperpolarization of neurons in layer2/3.

### Effect of multisensory stimulation on the individual neurons and the network

Given the large fraction of multisensory neurons within A1, we probed the nature of modulation during multisensory stimulus presentation. We took those neuron categories that showed significant modulation upon multisensory presentation as compared to unisensory presentation (MA, MAV, M0 & MV. Fig. 2, Table 1). We first compared the response properties of these neurons when presented with unisensory and multisensory stimuli and classified them as suppressed if the response to multisensory stimulus was significantly lower than the response to unisensory stimulus and enhanced if the response to multisensory stimulus was significantly greater than that to unisensory stimulus. The group of multisensory neurons whose responses were similar to both auditory and multisensory stimuli necessarily belonged to MAV (Methods, Classification of neurons).

Type MA neurons forms the greatest proportion of multisensory neurons in A1, equivalent to type A0 (MA=34%, A0=34%) (Fig. 2B, middle and Table. 1). Almost all of the MA cells (98%) showed significant suppression in responses upon multisensory presentation. Similar to A1, majority of MA neurons of secondary areas (AuD: 90%, AuV: 93%; Fig. 5A) also showed suppression upon multisensory presentation. Similarly for *MV* neurons, majority of which also undergo suppression in A1 and in AuV, while in AuD, 14.6 % neurons show enhancement of responses to multisensory stimulation. This percent can be explained by AuD’s proximity with secondary visual areas. AuD previously has been shown to be multisensory, responding to both auditory and visual stimuli (Sharma and Bandyopadhyay, 2020). Another point to notice here is that in the type MAV population, not all cells showed significant change on multisensory presentation (85%) and all those that did majority of them showed suppression (Fig.5A, Table 1). Both type MV and type MA population showed suppression of responses on multisensory presentation, regardless of the region specificity.

**Figure 5.**
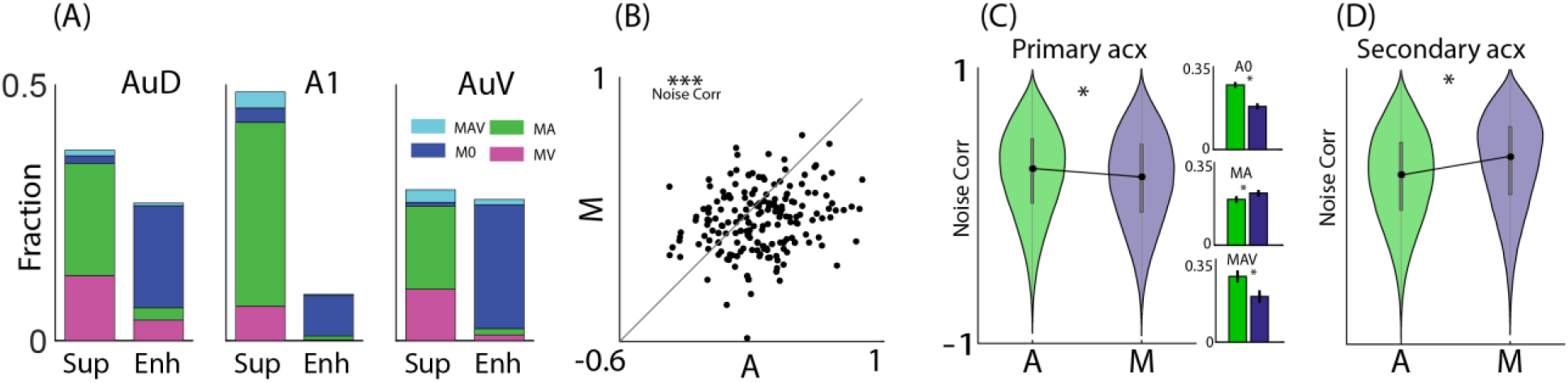
Multisensory modulation in the ACX. **A**, Multisensory neurons showing suppression and enhancement as a fraction of the total number of cells in that region, colour coded by type of neurons. Cyan, type MAV; blue, type M0; green, type MA; magenta, type MV neurons. **B**, noise correlations between pairs of simultaneously recorded ACX neurons during electrophysiology experiments (∗∗∗ p-value <0.001, paired t-test). **C** left, violin plot of population noise correlations in primary ACX during auditory (green) and multisensory (blue) presentation. The filled black dot shows the median of the population and the vertical grey line shows the first and third quartiles. ∗= significant based on 95% confidence interval from 1000 bootstraps. **C** right, bar plot of cell type wise mean noise correlations in the A1. Top (A0), middle (MA), bottom (MAV). Green (auditory), blue (multisensory). ∗= significant based on 95% confidence interval from 1000 bootstraps. Error bars show the bootstrap confidence interval. **D**, violin plot of population noise correlations in secondary ACX during auditory (green) and multisensory (blue) presentation.

Neurons with similar stimulus preferences have higher noise correlations (Ecker et al., 2011) and change in the correlations means change in the network involved. We attempted to decipher the network in auditory and multisensory stimulation based on noise correlations. We calculated the correlation coefficient on the mean subtracted trial data for each stimulus in the response window. We first looked at our electrophysiology data, recorded from deep layer 2/3 of the ACX, with auditory and multisensory stimulus (Fig. 5B, left). We observed that during multisensory presentation, there was a decrease in noise correlations among neurons (mean ± se: A; 0.26 ± 0.02, MS; 0.15 ± 0.02, paired t-test p<0.001, Fig.5B). We then computed the noise correlations during auditory and multisensory presentation from Calcium signals of auditory responding cell types (A0, MA, MAV) and corroborated it with the electrophysiology data. We found a similar trend from calcium signals as well, where there was a significant decrease (95% confidence with 1000 bootstraps) in noise correlations for multisensory stimulus (mean ± se A=0.24 ± 0.001, MS=0.19 ± 0.001, Fig.5C).

Given higher noise correlations for auditory stimulus as compared to multisensory, we looked at the contribution of auditory responding populations (A0, MA, MAV) to this effect (Fig.5C, right). We found that A0 (mean ± se A: 0.27 ± 0.002 MS: 0.18 ± 0.002, Fig.5C,top) and MAV (A: 0.27 ± 0.005, MS: 0.19 ± 0.005, Fig.5C, bottom) cells have significantly higher noise correlations (95% confidence interval on 1000 bootstraps) during auditory stimulation and low for multisensory stimulation. While MA cells (A: 0.19 ± 0.002, MS: 0.22 ± 0.002, Fig.5C, middle) show significantly higher noise correlations (95% confidence on 1000 bootstraps) for multisensory stimulus and low for auditory stimulation. MA neurons also have low noise correlations for auditory stimulus compared to that of A0 and MAV populations (Fig. 5C, right). Noise correlations can affect the neural population’s capacity to discriminate between two stimuli without affecting the responses of the individual neurons(Downer et al., 2015). This suggests that A0 population with very similar response profiles to auditory and multisensory stimulus relies on reduction in noise correlations for better discrimination of the two stimuli. Whereas MA cells with dissimilar response profiles appear to be less dependent on the population noise correlation for stimulus discrimination rather rely on response strengths to discriminate auditory and multisensory stimulus in A1. To compare the noise correlations obtained from secondary areas, AuD and AuV, we combined the pairs from A0, MA and MAV populations from both regions. Contrary to A1, secondary areas have significantly higher noise correlation (95% confidence for 1000 bootstraps) for multisensory stimulation compared to auditory (A: 0.21 ± 0.002, MS: 0.31 ± 0.002,Fig. 5D). This suggests that secondary areas do not rely on reduction in noise correlations for discriminability.

## Discussion

In this study, we show that neurons in the ACX have complex multisensory architecture, comprising of different neuron types that are affected by the presence of visual or multisensory stimulus in various ways. Contrary to previous reports, approx. 20% neurons in layer 2/3 of the ACX responded to visual stimulation, including neurons that only responded to visual (type V0 and MV) stimulus and others which responded to auditory stimulus as well (type MAV). This diversity of response types and thus of neurons in the ACX was unknown previously. However, our classification of V0 and MV neurons is based on the exercise that our auditory stimulus is a broadband noise. There could be a decrease in the number of these cells if a pure tone were to be used as the auditory stimulus. Nevertheless, these neurons would still be classified as visual responding multisensory cells.

When we look at the topography of the ACX at a greater resolution, we find neurons that respond to visual stimulus, in close proximity to the neurons that respond exclusively to the presentation of auditory stimulus (type A0). Type MV neurons which undergo modulation in presence of auditory stimulus were prevalent and are present within 100um vicinity to auditory specific neurons (type A0).

Multisensory stimulus do not modulate type A0 and type V0 neurons, they respond specifically to the features of sound and the presence of visual and auditory stimulus respectively does not affect their responses. These neurons may have a role in reliably transmitting information about auditory or visual stimulus irrespective of other stimuli in the environment, as they have reliable responses to their respective stimuli compared to other neuron types (Fig, 2C). Type MA and type MAV neurons are present within 50um radii around type A0 neurons (Fig. 3C). Type MA neurons were as common in primary ACX as were type A0 neurons, while they were greater in number in secondary areas. This close-knit architecture is indicative of high multisensory influence on the processing of auditory information and is capable of influencing auditory stimulus in multiple ways.

Response to visual stimulus differs between regions of the ACX. Greater numbers of neurons in secondary areas respond by excitation to visual stimulation compared to neurons in primary auditory area, where majority of neurons respond by hyperpolarization. This highlights a difference in processing of visual stimulus in primary and secondary areas and the possible role of primary ACX in processing visual information, which is to go silent. Silencing the primary sensory areas of non-sensory stimuli could aid in processing the sensory stimuli (Dehner et al., 2004), yet keeping the secondary areas concerning the non-sensory stimuli engaged to influence processing of the sensory stimuli. In addition, the local PV+ and SOM+ inhibitory neurons are not directly involved in causing these hyperpolarization responses but the likely source of this hyperpolarization is in the layer 5/6 of the ACX(Kapfer et al., 2007; Iurilli et al., 2012). Layer 5/6 of the ACX contains inhibitory neurons that do not respond to auditory stimulation but to visual stimulation. This is a novel finding and puts forward another role of feedback from deeper layers in influencing processing in the upper layers(Williamson and Polley, 2019). However, other studies on the role of auditory stimulus in processing visual information, shows a role of layer 1 mediated suppression in mice visual cortex (Iurilli et al., 2012) and role of thalamus mediated suppression in monkeys (Chou et al., 2020) influencing processing in layer 2/3 of the ACX. Similar processes can also exist in mice affecting processing in the ACX by the visual stimulus.

Although few, but there were certain neurons of type M0 which showed hyperpolarization on multisensory stimulation (Fig. 2A, upper row and left column). Occurrence of such responses should involve either summation of subthreshold inhibitory drive from neurons responding to auditory and visual stimulus separately or inhibitory neurons of type M0 responding with excitation. We, in fact, found inhibitory type M0 neurons responding with excitation to multisensory stimulus. These neurons could be a potential source of the hyperpolarization responses in type M0 population and a potential explanation of widely observed suppression on multisensory stimulation (Fig. 5A). Another point of difference between primary and secondary ACX stems from the noise correlation values during multisensory stimulation. There is an increase in noise correlations among the neurons in the secondary areas during multisensory stimulation while the correlations decrease in primary ACX. This decrease in noise correlations means an increase in discriminability between stimuli(Downer et al., 2015). Thus, primary ACX not only has a role in tone discriminability(O’sullivan et al., 2019) but also have network able to discriminate between cross-modal stimuli. Our classification of auditory cortical regions into A1, AuD and AuV is coarse(Kanold et al., 2014). Proper classification of different auditory cortical regions can yield more specific information and functions of the fields of the ACX. In addition, lack of temporal resolution in our recordings also limits our interpretations. Having access to greater temporal resolution could yield better circuit dissection, information about possible anatomical pathways as reported in the previous literature, and comparison of regions could be done in greater depth.

Several previous studies have shown the existence of visual inputs into the ACX, including humans(Calvert et al., 1999), non-human primates(Kayser and Logothetis, 2007) ferrets(Bizley and King, 2008b), gerbils(Hackett and Schroeder, 2009), rats(Wallace et al., 2004a). While most of them deals with modulation of auditory cortical activity during multisensory presentation, a limited number of these show spiking in the ACX by the visual stimulus in the deeper layers of the cortex (Morrill and Hasenstaub, 2018). Existence of these neurons in the superficial layers would allow tracking these neurons over days and establish the existence of these neurons more concretely. This will also allow us to selectively look at the role of a given multisensory population type and its role in affecting the processing of auditory and multisensory stimuli. Several previous studies have argued against the unimodal nature of primary cortices(Wallace et al., 2004b; Ghazanfar and Schroeder, 2006; King and Walker, 2012). This study is in concord with such studies and provides evidence for influence of multisensory stimulus on the auditory information processing.

